# Two matings lead to more copulatory wounding than a single mating in female *Drosophila melanogaster*

**DOI:** 10.1101/2025.02.20.639254

**Authors:** Bengisu S. Subasi, Anika L. Finsterbusch, Martha Büge, Sophie A.O. Armitage

## Abstract

Copulation can result in males inflicting wounds to the female genitalia, so-called traumatic mating. Such wounds are potentially costly as they could be entry points for infections, and they have been associated with a shorter lifespan in an insect species. In many species of insects, females mate with more than one male, which leads to the question of whether the number of matings affects the amount of genital damage that females suffer from those matings. Here we test whether copulation frequency affects the number or size of genital wounds in *Drosophila melanogaster*. Females who mated twice had more genital wounds and a larger total area of wounding, compared to females who mated once. However, females who refused to mate a second time had a similar area of wounding as females who mated twice. We found that wounds to the ventral abdomen also increased with increased mating frequency. Our results support the idea that polyandry can incur an under-appreciated cost in terms of increased female copulatory wounding. Investigating genital and abdominal wounds is crucial to a better understanding of the consequences of sexual conflict and the selective pressures shaping mating behaviour.

## 1. Introduction

The evolution of traumatic mating, i.e., the wounding of a mating partner by specialised reproductive structures during copulation (Lange *et al*. 2013), is one potential consequence of sexual conflict. Traumatic mating is phylogenetically widespread across animal taxa (Lange *et al*. 2013; Reinhardt, Anthes & Lange 2015). It is hypothesised to have evolved, for example, to allow male anchorage to the female during mating or to spur female reproductive investment (Lange *et al*. 2013). In insects, the resulting wounds are repaired by the immune system, which results in melanisation at the site of wounding (Whitten & Coates 2017). Melanised patches have therefore been used as indicators of wound repair in the genital tracts of several insect groups, e.g., beetles (Crudgington & Siva-Jothy 2000; Matsumura *et al*. 2017), ants (Baer & Boomsma 2006; Kamimura 2008), bedbugs (Bartonicka *et al*. 2023) and flies (Blanckenhorn *et al*. 2002; Kamimura 2007). Such copulatory wounds will be costly for the female, for example, if they provide an opportunity for microbes to enter the body (Siva-Jothy 2009; Otti 2015; Bellinvia *et al*. 2020), if they entail repair costs, or result in haemolymph loss (Reinhardt, Anthes & Lange 2015).

Polyandry is common across animal species (Taylor, Price & Wedell 2014), and it can benefit females by e.g., increasing reproductive output (Lee et al 2022) or by allowing for genetic bet-hedging (Garcia-Gonzalez et al 2015). However, there are also well-documented costs, and in insects these can include reduced hatching rate (Orsetti & Rutowski 2003) and reduced lifespan (Fowler & Partridge 1989; Crudgington & Siva-Jothy 2000; but see Yan *et al*. 2025). Copulatory wounding is almost certainly an additional cost of mating in general, for example female bean weevils with a higher number of genital scars had shorter lifespans (Gay *et al*. 2011). Although selection under polygamous conditions leads to males who scar females more (Gay *et al*. 2011), there is a paucity of studies that have quantified wounding in relation to a known number of matings (but see Bartonicka *et al*. 2023, who quantified the number of wounds to the spermalege, a paragenital organ of female bedbugs). Here, we hypothesised that copulatory wounding is an additional cost of polyandry, and we predicted that females mating with two males would receive more copulatory wounding compared to females mating with one male.

Female *D. melanogaster* typically mate with more than one male, and they are wounded during copulation. At least 23 species of the *D. melanogaster* species group have evolved traumatic mating (Kamimura 2010), including *D. melanogaster* (Kamimura 2007; Kamimura 2010). The traumatic mating strategy has since been ascribed to the sub-strategy of traumatic penetration, that is where copulatory wounding occurs without the direct injection of male-derived secretions (Lange *et al*. 2013). In *D. melanogaster* conformational changes of the male phallic parts occur during copulation, which results in genital wounding (Fig. 1a): The spiny ends of the left and right dorsal branches of the basal processes (more recently termed dorsal postgonites, Rice *et al*. 2019) penetrate the female reproductive tract (Kamimura 2010), and they have been hypothesised to play a role in anchoring the male to the female (Kamimura 2010). In *D. melanogaster* the usually twinned melanised patches indicative of wounding are found on or near the female lateral folds (Kamimura 2010; more recently termed the vaginal furcal dorsolateral fold, McQueen *et al*. 2022). Such genital damage has been found in wild-collected (Subasi *et al*. 2024) and lab-reared flies (Kamimura 2007). Furthermore, for more than a century it has been known that female *D. melanogaster* can mate with more than one male under lab conditions (Nonidez 1920), and that wild female *D. melanogaster* frequently mate with more than one male (Milkmann & Zeitler 1974; Ochando, Reyes & Ayala 1996; Harshman & Clark 1998; Imhof *et al*. 1998).

**Figure 1.**
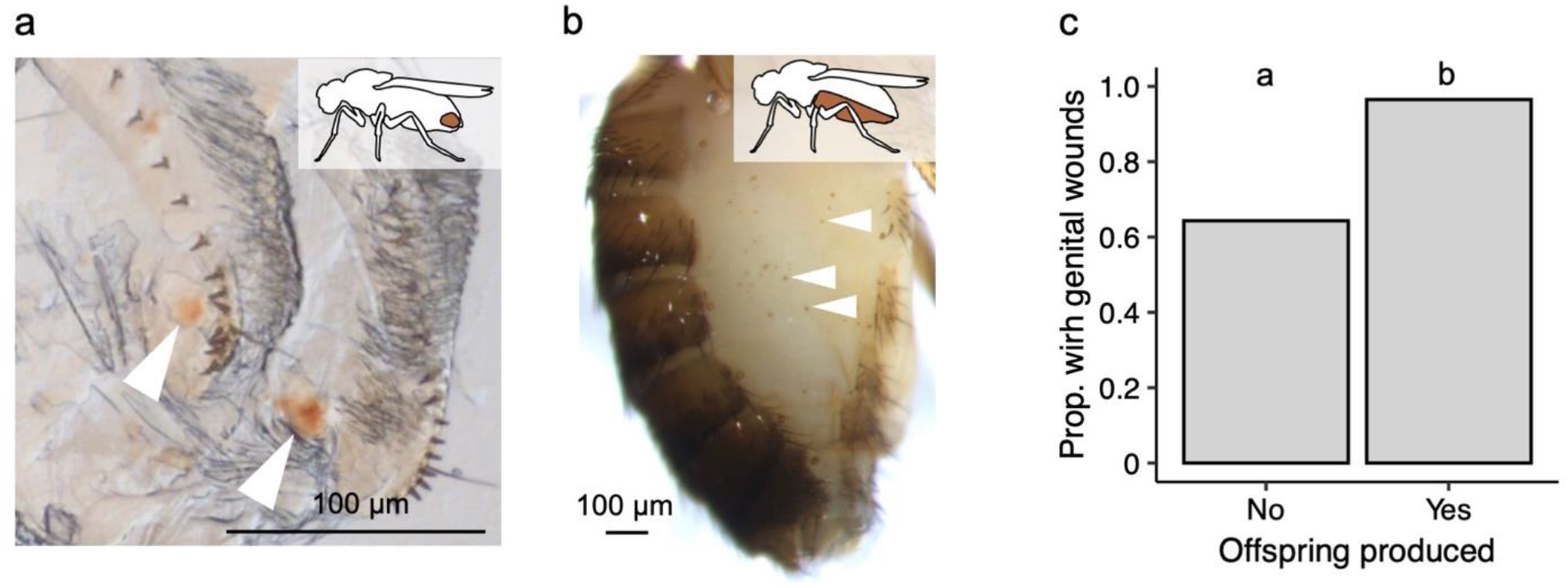
Genital and abdominal wounds, and wild-collected flies. (**a**) Genital wounding: white arrow heads point to mating-induced wounding on the female vaginal furcal dorsolateral fold and inset image indicates genital region. (**b**) Abdominal wounding: an example of abdominal wounds on a wild-collected female *D. melanogaster*. White arrowheads point to three of the spots, and inset image indicates the abdominal region checked for wounds. (**c**) The proportion of wild-collected females with wounded genital tracts in relation to whether they produced offspring or not after being brought into the lab. Sample sizes of all females who did not, and who did, produce offspring: n = 28 and 57 respectively. The different letters above the treatment groups indicate that the proportions are significantly different from each other.

Here we first investigated wounding prevalence in wild-collected *D. melanogaster* females, and its association with offspring production. We predicted that females producing offspring would be more likely to be wounded compared to females with no offspring. We then carried out two lab experiments to test whether copulation frequency affects the number or area of female *D. melanogaster* genital wounds. Harshman & Clark (1998) estimated that for 19 wild-inseminated females the mean number of males that mated with each female was 1.82, we therefore compared singly and doubly mated females. In our first experiment we predicted that females mated to two males would have more wounds and a larger total area of wounding compared to females mated to one male. There is genetic variation in the amount of harm (courtship and accessory gland proteins in seminal fluid) that male *D. melanogaster* inflict on females, which was measured as the impact on fecundity (Filice & Long 2016), therefore we used two *D. melanogaster* populations to test whether there is population-level variation in genital wounding. It was previously observed that wild-collected female *D. melanogaster* can have small melanised spots on their ventral abdomen (Subasi *et al*. 2024; Fig. 1b), which we hypothesised to be related to mating. To test this, we noted the occurrence of this abdominal wounding on mated and virgin females. Many females in the first experiment did not remate when given the opportunity. Therefore, in our second experiment we tested whether this self-selecting group of females had a similar degree of genital wounding to singly-mated females, or whether they showed intermediate or similar wounding to the doubly-mated females. To investigate the relationship between abdominal wounding and mating in more depth, we recorded the location and the number of abdominal wounds of all groups of females.

## 2. Material and methods

### (a) Traumatic mating in wild collected females

We examined wild-collected females for wounds in their genital tracts, and whether wounding relates to offspring production. Females morphologically identified as *D. melanogaster* and *D. simulans* were collected in early summer, late summer, and autumn of 2021, at three sites in and Berlin and Brandenburg, Germany (see supplementary information). After collection, the females were placed into individual food vials for six days, and we noted whether they produced offspring or not (presence of larvae), as we deemed the production of larvae as an indication that the mating had been successful. After six days the females were placed individually into microcentrifuge tubes containing 99% ethanol and stored at -20°C for later analysis. We collected 517 females in total, from which 432 produced offspring and 85 did not (see Table S1 for sample sizes according to collection site and season). Of these females, from across the collection seasons and sites, we randomly selected females who did and did not produce offspring (n = 58 and n = 47 respectively). We examined the genitalia of the male offspring to discriminate between *D. melanogaster* and *D. simulans*. Females who did not produce offspring were identified molecularly (see supplementary information) after they had been dissected to examine copulatory wounds. Female genitalia were dissected in October 2022 in a random order and blind with respect to sample collection place and time. The methods were the same as described for the below lab experiments, except we recorded whether wounding was present or absent, and not the number or area of wounds.

### (b) Experiment 1 – The effects of copulation frequency and population on genital wounding and the presence of abdominal wounding

Here we assessed the effects of copulation frequency (single or double mating) and population origin on the number and area of genital wounds, and on the presence of abdominal wounds. We used females who mated either once or twice and dissected the female genitalia to quantify wounding. We repeated the experiment described below three times. We note that it was previously suggested that *D. melanogaster* copulatory wounds do not show evidence of multiple matings (Kamimura 2010), although it was not mentioned how, or whether, wounding was quantified.

#### Fly populations and maintenance

We used two *D. melanogaster* populations. One was established in 2007 from 160 *Wolbachia*-infected fertilised females collected in Azeitão, Portugal (Martins *et al*. 2013) gifted to us by Élio Sucena, hereafter the PT population. The second population was established in 2021 from 402 fertilised females collected from three sites (see supplementary information) in Berlin and Brandenburg, Germany, hereafter the DE population. The populations were maintained in population cages containing at least 5000 flies, with non-overlapping generations of 14 days. We fed the flies with sugar yeast agar medium (SYA medium: 970ml water, 100 g brewer’s yeast, 50 g sugar, 15 g agar, 30 ml 10 % Nipagin solution and 3 ml propionic acid; (Bass *et al*. 2007) and maintained them at 25 °C, on a 12:12 h light-dark cycle, at 70 % relative humidity. The experimental flies were kept under the same conditions. For two generations prior to the start of each replicate experiment we reared the flies at a constant larval density of 100 larvae per food vial (95 x 25 mm) containing 7 mL of SYA medium (see supplementary information). Using CO_2_ for anaesthetisation, virgins were collected and placed into fresh food vials containing either 10 females or 20 males.

#### Experimental setup

In total we had four treatment groups (Table S2), and the subscript “d” refers to the number of days post adult eclosion: (i) 0_d5_; control (virgin) females who were not allowed to mate on day five, (ii) 0_d7_; control (virgin) females who were not allowed to mate on day seven, (iii) 1_d5_; single mated females who mated once on day five, and (iv) 2_d5 d7_; double mated females who mated once on day five and again on day seven. Females were randomly assigned to be control females or females given the opportunity to mate. In the afternoon before the first mating, four-day old adult virgin males were placed individually into food vials. Starting at nine o’clock the following morning, one person transferred five-day old females individually into vials containing a male from the same population as the female, alternating between setting up pairs from the two populations. Immediately after adding a female to a vial, we transferred it to an evenly lit observation shelf. A second person, who was blind to the identity of the population of origin, observed the vial to record the time of transfer, and the time the mating started and ended. From this data we calculated the latency to mate (time from transfer to start of mating) and the duration of mating. We gave each pair three-hours to start mating. Males were removed from the vials after copulation to prevent remating. At the same time as the mating females, we set up virgin females (0_d5_) which were each added to a food vial that did not contain a male (7 to 10 per population, per replicate). Between 62 and 84 females per population and replicate were randomly allocated to the double-mating group. Approximately forty-eight hours after their first mating, we transferred the double-mating group to food vials containing a virgin male, and the procedure was the same as for the first mating. Once again, we included 7 to 10 virgin females per population and replicate to be virgin controls (0_d7_) for the second mating. Twenty-four (± 1) hours after the end of the first or second copulation, the mated females were anaesthetised using CO_2_ and placed individually into 1.5 mL reaction vials, each containing 100 μl of 99 % ethanol. To examine whether the copulation had been successful, we kept the food vials that had contained the females in the 24-hours following copulation, and eight days later we recorded whether larvae were present.

#### Examination of genital wounds

One hour (± 30 mins) after being placed in ethanol, we removed the females and transferred them to a microscope slide with a drop of *Drosophila* Ringer’s solution (182 mmol KCl, 46 mol NaCl, 3 mmol CaCl2, 10 mmol Tris HCl; pH 7.2; (Werner *et al*. 2000) and placed them under a stereomicroscope (Leica M205C) at up to 160 × magnification. We applied pressure to the female abdomen so that the terminalia was extruded, a second pair of forceps was used to gently pull the terminalia away from the abdomen, and the terminalia was placed onto another microscope slide with 5 μl of Ringer’s solution. A cover slip was placed on top to flatten out the terminalia to examine the vaginal furcal dorsolateral folds. The person observing melanised spots and dissecting the flies was blind to the identity of the flies and their treatment, and they were processed in a randomised order. Between 6 and 16 females per experimental replicate and population were successfully dissected and photographed for the single mating group, and the number was between 3 and 13 for the double mating group.

We visualised the dissected genitalia using a light microscope (ZEISS Germany Axiophot) and one photograph was taken of each tract using a Jenoptik ProgRes CF digital camera, at a total magnification of x 250. The photographs were analysed using ImageJ (Rasband 1997-2018) version 1.53a. Wounds appear as brown spots on the genital tract (Kamimura 2007, and see Fig. 1a). We recorded the number of wounds and their total area on one side of the vaginal furcal dorsolateral folds, as in most photographs only one side was visible. In preliminary tests, we did not find significant differences between the average of measuring two sides compared to the measurement from one side only. We traced the outline of each wound and the total area per female was recorded in μm^2^. The repeatability of measuring the area of the wounds with the above analysis methods was high (see supplementary information). A preliminary experiment showed that the area of the wound did not change significantly if females were sacrificed one, three or six days after mating (Fig. S1).

#### Examination of abdominal wounds

A previous study showed that wild-collected females exhibit more melanised spots on their ventral abdomen compared to males, and that similar wounds are absent on the female dorsal abdomen and in males (Subasi *et al*. 2024). Based on these findings, we focused exclusively on the ventral abdomen to test whether there is an association between melanised spots and mating. After mating and prior to genital dissection, females were placed in a drop of *Drosophila* Ringer’s solution and examined under a stereomicroscope (Leica M205C) at up to 160× magnification. We carefully inspected the ventral abdomen of mated and virgin females and recorded the presence/absence of melanised spots in experimental replicates two and three.

### (c) Experiment 2 – The effects of single matings, double matings, and not remating, on genital and abdominal wounding

The first set of experiments resulted in a group of self-selecting females who did not mate a second time, and in this second experiment we asked whether this group were wounded to a similar degree to the singly or doubly mated females, or whether they had an intermediate amount of wounding. Our second aim in this experiment was to examine abdominal wounding in relation to mating, by recording the location and the number of abdominal wounds in all female treatment groups. We used only the PT *D. melanogaster* population, and the maintenance of the flies and experimental procedures followed the same methods as described above. We repeated the experiment described below three times.

In addition to the previously established treatment groups in experiment 1, two additional groups were included (Table S3): (v) 1_d7_; females mated once on day seven to control for possible differences associated with female aging and (vi) 1_d5, no d7_, females mated on day five but who did not remate on day seven despite being exposed to a male, i.e., the self-selecting group of females who did not remate. Ten females per replicate were randomly allocated to the virgin control group (0_d7_). Per experimental replicate, between 176 to 379 females were randomly allocated to single mating treatments (1_d5_ and 1_d7_), while between 112 to 187 females were randomly allocated to the double mating treatment (2_d5 d7_). Of the females who refused to mate a second time, 15 were randomly allocated to the group that mated on day five but did not remate on day seven (1_d5, no d7_).

To determine whether melanised spots on the ventral abdomen were caused by mating, flies in some treatment groups were examined both before and after mating (see Table S3). Unlike experiment 1, females were examined for abdominal wounds prior to the first mating. On the day before the first mating, four-day-old adult virgin females were briefly anesthetized with CO2 and assessed for abdominal wounds. To do this, females were placed on a small piece of parafilm, and a drop of water was applied to their abdomen to enhance the visibility of melanized spots. Preliminary tests confirmed that the water did not influence melanized spot formation. Each female was examined under a stereomicroscope (Leica M205C) at up to 160× magnification. We noted on which of tergite numbers 1 to 7 the melanised spots were located, and how many spots each female had. After this the females were placed individually into numbered vials containing fresh food.

The following day, mating was carried out as described in experiment 1. Twenty-four hours after the first mating, 0_d5_ and 1_d5_ females were again examined for abdominal wounding. After examining the abdomen, mated females were dissected to quantify copulatory wounds as described above. Before the mating on day 7, females from the 0_d7_ and 1_d7_ groups were examined for abdominal wounds. After the second mating opportunity, females from the 1_d5, no d7_ and 2_d5 d7_ were examined for melanised spots, and dissections were performed to assess copulatory wounds.

### (d) Statistical analyses

To perform the analyses, R (R Core Team 2023) version 4.2.2 and RStudio (Posit Team 2023) version 2024.09.0+365 were used. Details of all models are given in the supplementary information.

## 3. Results

### (a) Wild collected females show genital wounding

Eighty one percent (n = 85) of the randomly selected wild-collected females were *D. melanogaster*. Of the *D. melanogaster* females, 86 % had genital wounds. Wild-collected females who produced offspring in the lab were more likely to be wounded than females who did not produce offspring (Chi square = 11.03, df = 1, p = 0.0009; Fig. 1c). Indeed 97 % of females who produced offspring had visible wounds, although we note that a substantial proportion of females who did not produce offspring in the lab also had visible wounds (Fig. 1c).

### (b) Lab experiments

#### Proportions of females mating at the first and second mating attempts

In the first experiment, 81.3 % ± 2.95 (SE) of females copulated at the first mating opportunity, and around 13 % ± 2.43 (SE) of females mated a second time. There was no effect of population on the frequency of females that mated at the first (Chi square = 3.70, df = 1, p = 0.055) or second mating (Chi square = 0.20, df = 1, p = 0.658). In the second experiment, 62.3 % ± 5.19 (SE) of females copulated at the first mating opportunity whereas only 3.6 % ± 1.27 (SE) of females mated a second time.

#### Latency to mate was longer for the second mating

Latency to mate was significantly longer for the second mating compared to the first mating in both experiments 1 and 2 (Table S4; Fig. S2a & b). In experiment 1, the DE population started mating more quickly than the PT population (Table S4). In contrast to latency, in experiment 1 copulation duration was marginally longer for the first mating compared to the second mating, but it did not differ in experiment 2 (Table S4; Fig. S2c & d).

#### Genital wounding is affected by mating frequency

As expected, none of the virgin females (n_exp1_ = 114) but all of the mated females (n_exp1_ = 200) had visible genital wounds. In the subset of females who mated once, for both experiments, there was no effect of latency to mate or copulation duration on the number of wounds, or the area of wounding (Table S5).

As predicted, across both experiments females who mated twice received more wounds compared to singly mated females (Table 1; Fig. 2a & b). Furthermore, the females who did not remate (1_d5, no d7_), had a similar number of wounds to both single and double mated females (Table S6; Fig. 2b). Wound area showed a similar effect of mating frequency as wound number, i.e., the wound area was larger when females mated twice compared to once (Table S6; Fig. 2c & d). However, in contrast to wound number, wound area of the females who did not remate (1_d5, no d7_), was significantly larger than singly mated females, but similar to the females who mated twice (Table S6; Fig. 2d). In experiment 1, neither wound number nor wound area were affected by the fly population or an interaction between mating treatment and fly population (Table 1). Across both experiments there was a positive relationship between the total number of wounds and the area of wounding (Table 1; Fig. 2e & f).

**Table 1.**
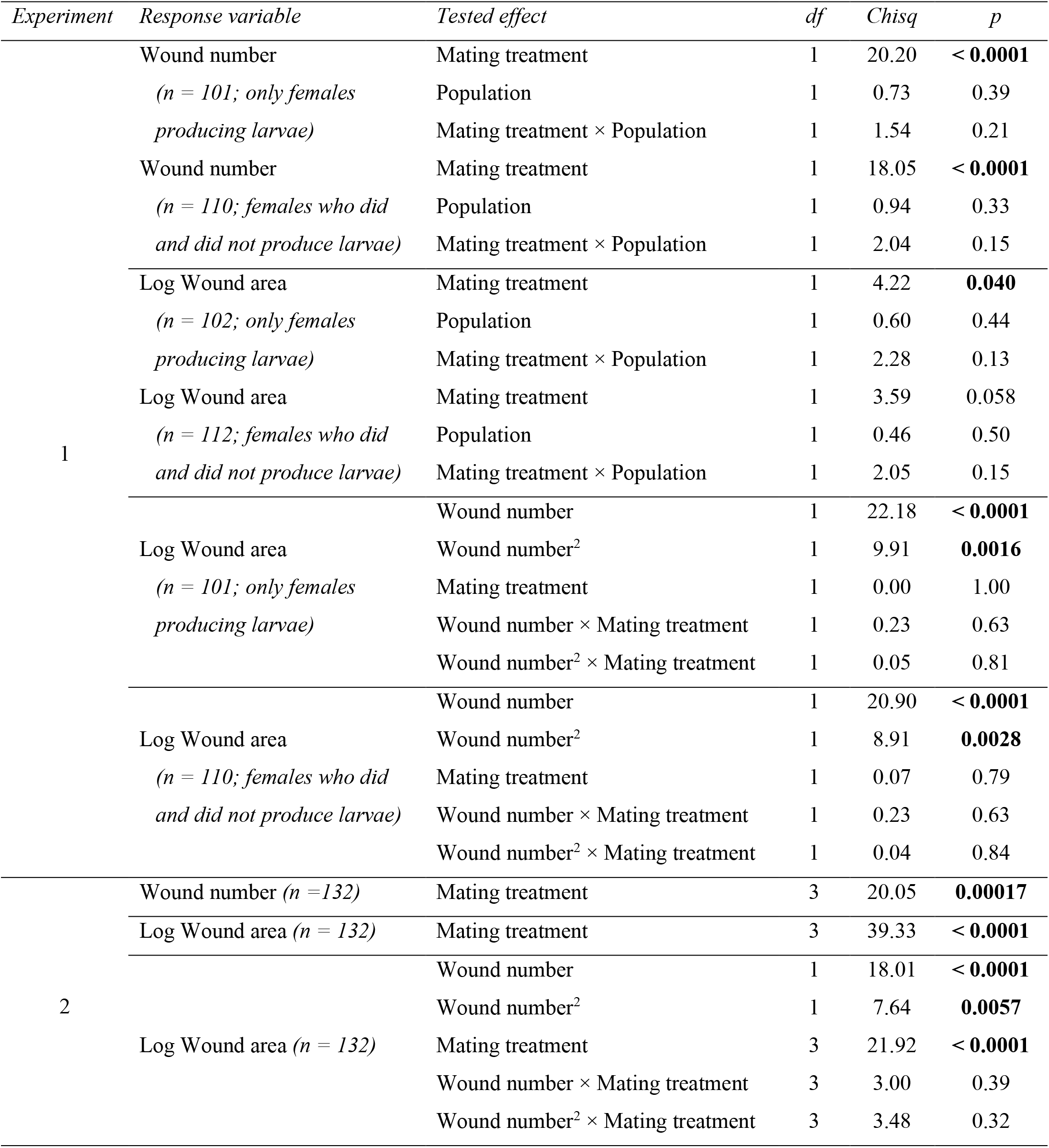
The effects of mating frequency and population on genital wound number and area. Each response variable was first tested using only females who produced larvae after the first mating, and then using all females, i.e., also females who did not produce larvae after the first mating. Sample sizes are given in the table.

**Figure 2.**
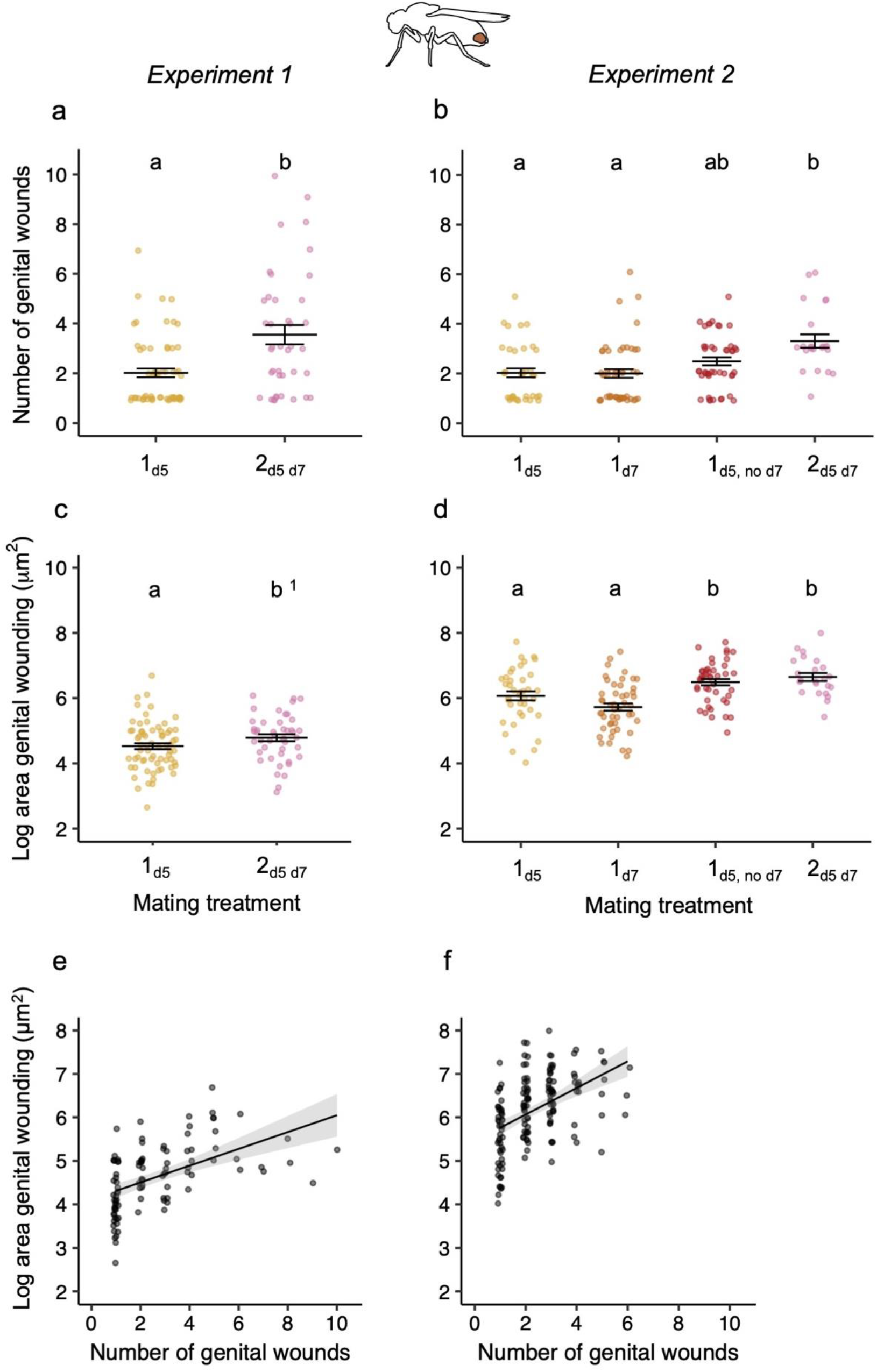
The effect of the number of matings on genital wound number and area. The effect of the number of matings on the number of wounds observed for (**a**) experiment 1 and (**b**) experiment 2, and on the area of wounding for (**c**) experiment 1 and (**d**) experiment 2. The relationship between the number and area of genital wounds for (**e**) experiment 1 and (**f**) experiment 2. Experiment 1 sample sizes for the number of wounds: 1_d5_ = 61 and 2_d5 d7_ = 40, and for area: 1_d5_ = 61, 2_d5 d7_ = 41. Experiment 2 sample sizes for both variables: 1_d5_ = 39, 1_d7_ = 49, 1_d5, no d7_ = 45 and 2_d5 d7 =_ 23. Only females who mated for a minimum of 10 mins and who produced larvae after the first mating are included. Each data point is from one female and data points are jittered for the number of wounds and the treatment groups. Means and standard errors are shown. When letters above the treatment groups are not the same, the treatment groups are significantly different from each other. ^1^When the area of wounding was analysed including females who did not produce larvae, p = 0.058, see Table 1.

#### Abdominal wounding is affected by mating frequency

In experiment 1, the presence of abdominal wounds on the ventral abdomen was almost completely related to mating treatment: less than one percent of virgins had visible abdominal wounds whereas more than 95 % of mated females had visible abdominal wounds (Fig. 3a; Table S7). Post-hoc comparisons showed that females that mated once or twice were more likely to have abdominal wounds than virgin females (Table S8).

**Figure 3.**
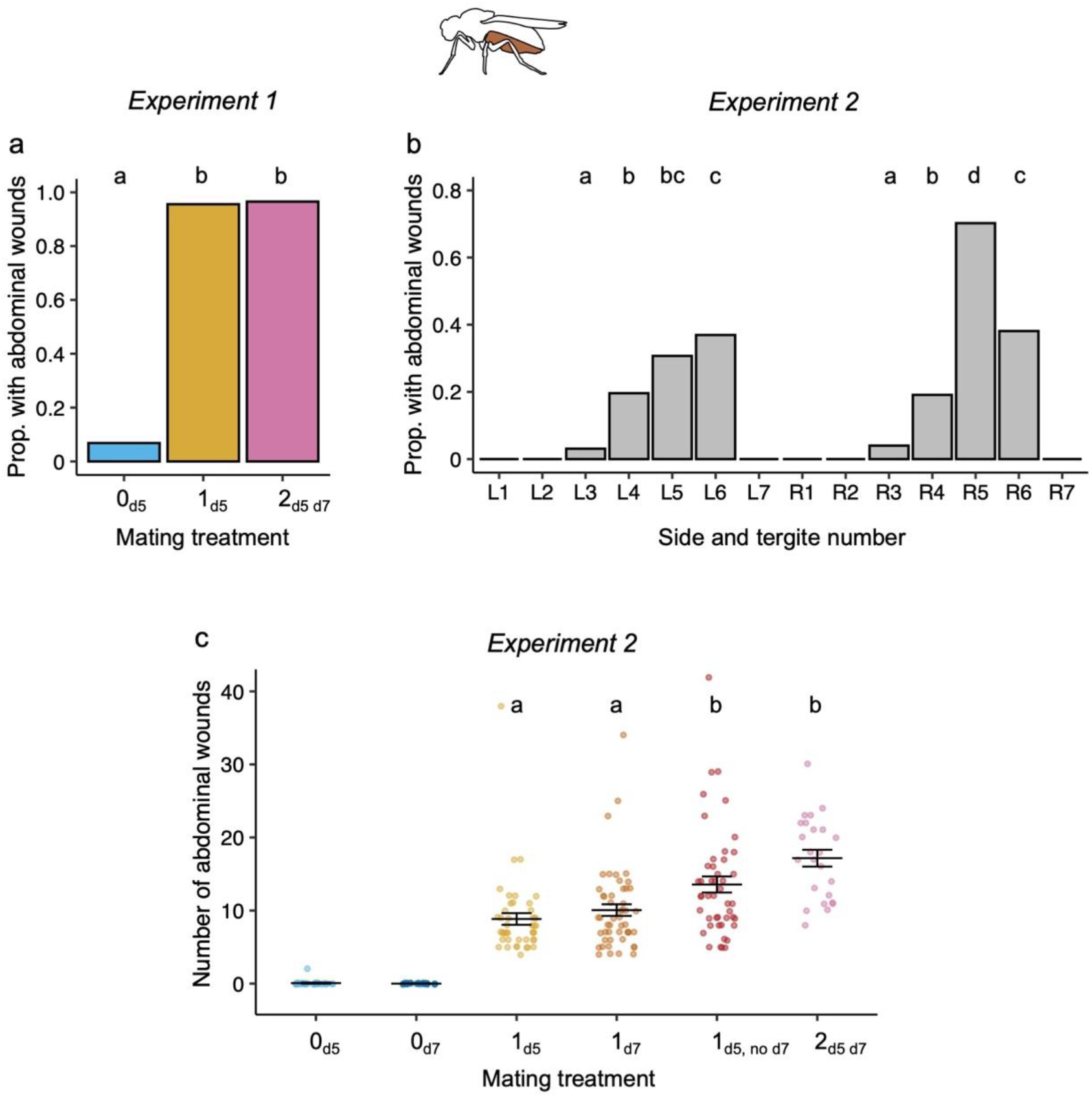
The location of abdominal wounds, and the effect of the number of matings on abdominal wound frequency. Experiment 1: (**a**) The proportion of virgins (0) and females mating once (1) or twice (2) who had melanised spots visible on the exterior of their ventral abdomen. Sample sizes of all females in each treatment group: 0_d5_ = 73, 1 _d5_ = 68, 2_d5 d7_ = 29. Here and elsewhere, when letters above the treatment groups are not the same, the treatment groups are significantly different from each other. Experiment 2: (**b**) The proportion of females who had abdominal wounds on their tergites (1-7, where 1 refers to the tergite closest to the thorax), and whether the spots were on the left- (L) or the right- (R) hand side of the tergites. The total sample size is n = 235. Note that treatment groups with proportions of zero (L1, L2, L7, R1, R2 & R7) were not included in post-hoc multiple testing. (**c**) The number of ventral abdominal spots found on each female for each of the mating treatments. Sample sizes are: 0_d5_ = 30, 0_d7_ = 30, 1_d5 =_ 45, 1_d7_ = 50, 1_d5, no d7_ = 46 and 2_d5 d7_ = 24. The two virgin groups were not included in post-hoc multiple testing.

In experiment 2, we investigated the location and number of abdominal wounds on the ventral abdomen. The abdominal tergites varied significantly in the number of wounds they received following mating (Chi-square = 269.13, df = 7, p < 0.0001). The wounds were found exclusively on tergites three to six, with 70 % of females showing wounds on the fifth tergite on the right-hand side of the abdomen (Fig. 3b). Consistent with experiment 1, abdominal wounds were strongly associated with mating treatment. Only one virgin female had two abdominal wounds before mating, and this number remained unchanged after mating (Fig. 3c). No singly mated females had wounds before mating and all females had at least four wounds on their abdomen after mating (Fig. 3c). Overall, there was a significant effect of mating frequency on the number of wounds (Table S6). Similar to the area of genital wounds, females that mated twice had more abdominal wounds compared to those mated once (Fig. 3c; Table S8), and the number of wounds on the females who did not remate (1_d5, no d7_), was significantly higher than singly mated females, and did not differ from the females who mated twice (Fig. 3c; Table S8).

#### No relationship between the number of abdominal wounds and genital wounding

There was no relationship between the number of abdominal wounds and either the number (Fig. S3a; Table S9) or the area of genital wounds (Fig. S3b; Table S9).

## 4. Discussion

Here we show that almost all wild-collected females who produced offspring had visible genital wounding. In our lab experiments, as predicted, two successful matings resulted in more wounds and a larger total area of wounding to the female genitalia, compared to when females were only allowed to mate once. We furthermore identified ventral abdominal wounding as being almost exclusively found in mated females. Abdominal wounds showed a similar pattern to genital wound area: females that mated twice had more abdominal wounds than those mated once. Therefore, in *D. melanogaster*, increased copulatory wounding is an additional cost of polyandry.

A previous study of wild-collected females showed that 98.4 % had sperm in their reproductive tracts, i.e., had mated (Markow, Beall & Castrezana 2012). Consistent with this number, here we show that of the females who produced larvae, 97 % were wounded. However, we also found that many wild-collected females had genital wounds but did not produce offspring (64 %). Given that the results from the lab show that virgins do not have wounds (this study, Kamimura 2007) it seems unlikely that the 64 % of females were unmated, instead it seems more probable that the conditions in the lab were not conducive for producing offspring, or that females had exhausted their sperm supplies.

Traumatic mating is one expression of sexual conflict, for example a single mating with a male seed beetle, *Callosobruchus maculatus*, evolved under conditions with increased sexual conflict (polygamy), resulted in more female copulatory damage compared to mating with a male evolved under monogamous conditions (Gay *et al*. 2011). We here show that females mating twice had more wounds, and a greater area of wounds compared to females that mated once. We are only aware of two other studies that have tested whether female genital wounding is affected by mating frequency when mating number is controlled: Blanckenhorn *et al*. (2002) found that the proportion of female dung flies, *Sepsis cynipsea*, with genital wounds was higher after two copulations (∼30 %) compared to after one copulation (10 %), and Bartonicka *et al*. (2023) found a positive relationship between the number of matings and the number of scars to the paragenital organ of female common bedbugs, *Cimex lectularius*. The increase in proportion or number of wounds with increasing mating number is consistent with our findings when comparing treatments 1_d5_ and 1_d7_ with the 2_d5 d7_ treatment. However, in *D. melanogaster* every mating results in at least one scar (Kamimura 2007; the present study), whereas not every dung fly copulation resulted in wounding (Blanckenhorn *et al*. 2002), and on average 3.5 matings led to one bedbug scar (Bartonicka *et al*. 2023). As might be predicted, and similar to Gay *et al*. (2011), we found a positive correlation between wound area and the number of genital wounds.

The self-selecting group of females who refused to remate (1_d5, no d7_) showed an intermediate number of matings between the single and double mated females, and a similar area of mating to the double mated females and more than the singly mated females. We reasoned that the genital wounding in this group resulted from the first copulation, as these females did not copulate successfully a second time. One hypothesis to arise from these findings is that the increased area of genital wounding in this self-selecting group of females compared to singly mated females is linked to the refusal to remate, for example if larger wounds are more costly to repair and if this affects the remating rate.

It was previously documented that wild-collected females have more wounds on the ventral abdomen compared to males (Subasi *et al*. 2024). Here we show that the wounds were predominantly located between the third and sixth segments on the ventral abdomen, i.e. on the terminal half of the abdomen, under which the female reproductive organs lie. We found that these abdominal wounds are associated with mating: less than one percent of virgins, but almost all (> 95 %) mated females had visible spots. In line with our findings for genital wounding, females mating twice had more abdominal wounds compared to females that mated once. Furthermore, females that mated once but refused a second mating showed a similar number of abdominal wounds to those that mated twice. However, despite the similar responses of genital and abdominal wounding to mating frequency, we did not detect a positive relationship between the number or area of genital wounding and the number of abdominal wounds, which might suggest that these two areas of wounding are somewhat independent from each other.

Unlike the genital wounds, the abdominal wounds are visible on the exterior of the ventral abdomen. It was previously hypothesised that the ventral abdominal wounds could be due to external damage caused by males during copulation (Subasi *et al*. 2024). In *D. melanogaster* males exhibit a series of courtship behaviours including tapping, following, wing vibrations, licking, abdominal bending, lunging, and grasping of the female (Spieth 1952; Hall 1994). During courtship it has also been reported that the tarsi of the males forelegs, are extended under the abdomen of the female and vibrate against it (Spieth 1966), or that the forelegs are used to grasp the abdomen (Cook 1975). It is unknown whether any of these behaviours would be able to cause the abdominal wounds that we observed. A non-mutually exclusive alternative explanation is that the abdominal wounds could be due to an internal physiological response to copulation itself (Subasi *et al*. 2024), which may originate from autoimmune damage elicited by mating or substances/pathogens (e.g., disseminated melanisation after pathogen injection, see Ayres & Schneider 2008) transferred at mating. Future studies will be needed to uncover the exact mechanisms.

Our experiment also allowed us to examine the proportion of remating females. Previous lab studies have shown that female remating rates can vary considerably according to exposure time to males (Fuerst, Pendlebury & Kidwell 1973), remating interval (Smith *et al*. 2017; Yan *et al*. 2025) and genotype (Chow, Wolfner & Clark 2013). Chow et al (2013) found remating rates between ∼10% and 100% for the DGRP fly lines that they tested, therefore the remating rates in the present study are at the bottom end of that distribution, however there are differences in the experimental set up used by that study and our own. One potential explanation for the lower remating rates in our second experiment compared to the first experiment could be the fact that the females were anaesthetised with CO_2_ more frequently in the second experiment because of the checks for abdominal wounds, and this form of anesthetisation can increase copulation latency (Barron 2000). Consistent with Pavković-Lučić & Kekić (2009), we found that latency to mate was longer for the second mating compared to the first mating. Copulation duration was longer for the first mating in experiment 1, which is consistent with some studies (Pavkovic-Lucic & Kekic 2009; Sirot, Wolfner & Wigby 2011) but not with other studies (Singh & Singh 2004; Friberg 2006; Lüpold *et al*. 2011).

In conclusion, our findings support the idea that multiple mating incurs a potential cost in terms of increased genital and abdominal wounding in female *D. melanogaster*, potentially representing a trade-off for females seeking benefits (e.g., genetic diversity) from multiple matings. However, females who refused to remate incurred similar damage to twice mated females, highlighting the importance of investigating this self-selecting group of females: whether or not the higher amount of genital wounding in this group is causally related to the lack of remating remains to be explored. In *D. melanogaster* systemic infections can result when females mate with males carrying pathogenic bacteria on their genitalia (Miest & Bloch-Qazi 2008). The mechanism for the latter is untested, but one possibility is that bacteria pass through copulatory wounds into the haemocoel, as was demonstrated for microbeads in *Drosophila santomea* and *Drosophila yakuba* (albeit at low rates, Kamimura 2012): If this is the case, more numerous or larger total wounds after multiple mating could further increase infection risk.

## Ethics statement

This work did not require ethical approval from a human subject or animal welfare committee. We obtained permission from landowners and managers to conduct the fieldwork detailed in this study. No other fieldwork permissions were required.

## Data availability

The datasets generated in this study and the R script containing statistical analyses will be available on Dryad.

## Competing interests

We have no competing interests.

## Acknowledgements

We thank Diana Aldana Alvarez, Lukas Maas, Malavika Menon and Laura Lender for their technical assistance in the lab, Baybora Gündüz for preliminary experiments, Mathias Franz for statistical advice, Oliver Otti for providing feedback on an earlier draft of the manuscript, and Mrs. and Mr. Lindicke, Jens Scheidereit and Annika Siegel for allowing us to sample on the farms.

## Funding

This study was supported by a Heisenberg Fellowship (AR 872/4-1 and AR 872/7-1) to S.A.O.A. from the Deutsche Forschungsgemeinschaft (DFG; https://www.dfg.de/en/index.jsp).

B.S.S. was supported by the DFG through project AR 872/5-1, awarded to S.A.O.A. as part of the Research Unit FOR 5026 “InsectInfect”.

## Authors’ contributions

B.S.S.: investigation, methodology, supervision, validation, visualisation, writing – review and editing

A.L.F.: investigation, methodology, validation, visualisation, writing – review and editing

M.B.: investigation, methodology, validation, writing – review and editing

S.A.O.A.: conceptualisation, data curation, formal analysis, funding acquisition, methodology, project administration, resources, supervision, visualisation, writing – original draft, writing – review and editing.

## Notes

### Competing Interest Statement

The authors have declared no competing interest.

