## Supplementary file for "Two matings lead to more copulatory wounding than a single mating in female *Drosophila melanogaster*"

This document includes:
Supplementary methods
Tables S1-S9
Figures S1-S3
References for supplementary information

### Supplementary methods

#### (a) Traumatic mating in wild collected females

Wild *Drosophila melanogaster* and *Drosophila simulans* were collected from three sites in Berlin and Brandenburg: Domäne Dahlem (denoted as D, 52.45883°N, 13.28901°E), Obsthof Lindicke (denoted as L, 52.38012°N, 12.86828°E) and SL Gartenbau (denoted as G, 52.69889°N, 13.08673°E). For molecular species identification, we extracted DNA from the legs, head, or whole body of each wild-collected fly. As a control for each species, we removed legs from individuals originating from lab-reared populations of *D. melanogaster* and *D. simulans*. We followed a modified DNA extraction protocol based on Gloor et al. (1992), where the modifications meant homogenising the samples with a 3 mm tungsten bead (QIAGEN) using a Retsch MM400 for three minutes at 20 Hz. After incubation, we centrifuged the mixture for one minute at 15,000 rcf and transferred the resulting supernatant to a new microcentrifuge tube. A PCR was carried out using the *Slif* primers, as described by Faria & Sucena (Faria & Sucena 2017), except that we used KAPA HiFi HotStart ReadyMix PCR kit (Roche).

#### (b) Lab experiments 1 and 2

For constant larval density, grape juice agar plates (25 g agar, 300 mL red grape juice, 21 mL Nipagin [10% solution] and 550 mL water (Koppik & Fricke 2017) were smeared with a thin layer of yeast and put overnight into the population cages. The following morning the plates were removed and placed into the incubator for 24 hours. After this, 100 first instar larvae were picked from the plates and transferred into each food vial.

To test the repeatability of measuring the area of the wounds, the photographs from 35 females from a separate experiment were analysed twice on two different days, and the repeatability was calculated according to Lessells and Boag (Lessells & Boag 1987). The repeatability of measuring the area (intraclass correlation coefficient, $r = 0.98$, $F_{34,35} = 2180$, $p < 0.0001$) of the genital tract twice from the same photograph was high. The repeated measurements were also correlated highly with each other (area: $rho = 0.99$, $S = 22.5$, $p < 0.0001$).

**(c) Statistical analyses**

For residual diagnostics of the statistical models, the DHARMa package (Hartig 2022) was used. The car package (Fox & Weisberg 2019) was employed for type II ANOVAs. Data visualization was carried out using ggplot2 (Wickham 2016). We also used the following packages for visualisation and data summaries: dplyr (Wickham *et al.* 2022), PairedData (Champely 2018), plyr (Wickham 2011), purrr (Henry & Wickham 2020), plotrix (Lemon 2006), qqplotr (Almeida, Loy & Hofmann 2018) and tidyr (Wickham & Girlich 2022). Experimental replicate was included as a random factor in each model, unless stated otherwise. We fitted generalised linear mixed models (glmms) using the glmmTMB package (Brooks *et al.* 2017) and lme4 (Bates *et al.* 2015). Post-hoc multiple comparisons were performed with the fifer (Fife 2014) (Fife, 2014) or emmeans (Lenth 2024) packages, using the Bonferroni adjustment for multiple comparisons.

*Wild-collected females*

To assess the effect of offspring production on the presence of wounding, we fitted a model with a binomial distribution, with presence of wounding included as a binary response variable.

m00: copulatory wound ~ offspring

*Lab experiments 1 and 2*

In experiment 1, we first asked if the propensity to mate at either the first or the second mating was affected by the fly population. Separate models were run for the first and second mating, and both used the family = "binomial" argument. The binary response variable utilised 1 for females that had mated and 0 when they had not mated:

m0a: mating success$_{\text{first mating}}$ ~ population + (1|experiment)

m0b: mating success$_{\text{second mating}}$ ~ population + (1|experiment)

In experiments 1 and 2, we created a subset of flies consisting of the females who had mated twice (n = 56 and n = 22, respectively). To test whether latency to mate or copulation duration differed for the first and second matings, we fitted models with either log transformed latency to mate, or mating duration as response variables. In experiment 1, the factors were whether it was the first or the second mating, and the population, as well as the interaction between the two. In experiment 2, we only used one population, so the only factor included in the models was whether it was the first or the second mating. To solve a convergence problem with model m1b.2, experiment” was removed from the model as a random effect. To account for the repeated measure from each female, female identity was included as a random effect. To allow for a random slope, female identity was nested within “first or second mating”:

Experiment 1 models:

m1a: natural log (latency to mate) ~ first or second mating × population + (1|experiment) + (first or second mating|female ID)

m1b: mating duration ~ first or second mating × population + (1|experiment) + (first or second mating|female ID)

Experiment 2 models:

m1a.2: natural log (latency to mate) ~ first or second mating + (1|experiment) + (first or second mating|female ID)

m1b.2: mating duration ~ first or second mating + (first or second mating|female ID)

Kamimura (2010) found that genital coupling was established after 10 minutes in all copulating pairs that were studied. Therefore, for further analyses we excluded all copulations that lasted for less than 10 minutes. This resulted in excluding four females for the first mating, and two for the second mating for experiment 1. In experiment 2, only one female was excluded for the first mating. We deemed the production of larvae as an indication that the mating had been successful. However, some females for whom we had mating data did not produce larvae after the first mating, so we therefore analysed the data both with and without females that produced larvae. In experiment 1, five females did not produce larvae in the wound number dataset, and six females did not produce larvae in the wound area subset. In experiment 2, all the females produced larvae, therefore we analysed the data only with the females with larvae.

In both experiments 1 and 2, we tested whether there was any effect of latency to mate or copulation duration on genital wounding after one mating. We fitted models with response variables as the number of wounds (family = poisson) and the natural log of wound area (family = gaussian). For the model, we applied a transformation by subtracting 1 from the number of wounds. Latency and copulation duration were included as covariates:

m2a: (wound number -1) ~ latency + duration + (1|experiment) m2b: natural log (wound area) ~ latency + duration + (1|experiment)

In experiment 1, we then tested whether there was any effect of mating treatment (one or two matings), population or their interaction, on genital wounding. In experiment 2, we focused only on the effect of mating treatment on genital wounding. Additionally, in both experiments, we tested whether there was a relationship between wound number and area. We fitted models with response variables as the number of wounds (family = poisson) using a transformation by subtracting 1 and the natural log of wound area (family = gaussian). For these analyses, in experiment 1, there were nine females who had not produced larvae in the wound number dataset and ten females who had not produced larvae in the wound area subset:

Experiment 1 models:

m3a: number of wounds ~ mating treatment × population + (1|experiment)

m3b: natural log (wound area) ~ mating treatment × population + (1|experiment)

m3c: natural log (wound area) ~ (number of wounds^2) × mating treatment + number of wounds × mating treatment + (1|experiment)

Experiment 2 models:

m3a.2: (number of wounds -1) ~ mating treatment + (1|experiment)

m3b.2: natural log (wound area) ~ mating treatment + (1|experiment)

m3c.2: natural log (wound area) ~ (number of wounds^2) × mating treatment + number of wounds × mating treatment + (1|experiment)

For experiment 1, to test whether the presence of abdominal wounds was associated with mating, we fitted a model with presence of black spots on the abdomen as a binary response variable (family = binomial). The factors were the mating treatment (virgin, one- or two-matings) and the population.

m4a: abdomen wound ~ mating treatment + population + (1|experiment)

In Experiment 2, we tested whether the location of the wounds on the ventral abdomen varied across abdominal tergites. To do this, we performed a Chi-square test using the number of the females with and without wounds on tergites three to six. Since no wounds were observed on the first, second and last segments, these were excluded from the analysis. Lastly, we tested whether mating treatment has an effect on the number of abdominal wounds. To do this, we fitted a generalised linear model with negative binomial distribution, where the number of abdominal wounds was the response variable and mating treatment was the predictor.

m4b: number of abdominal wounds ~ mating treatment + (1|experiment)

For experiment 2, we tested whether there is a relationship between the number of abdominal wounds and the number of genital wounds. To do this, we fitted a generalised linear model with the number of genital wounds as the response variable and the number of abdominal wounds as the predictor (family = gaussian).

m5a: number of genital wounds ~ number of abdominal wounds + (1|experiment)

Additionally, we tested the relationship between the number of abdominal wounds and genital wound area with using a generalised linear model, where the natural log of wound area was the response variable and the number of abdominal wounds the predictor  (family = gaussian).

m5b: natural log (wound area) ~ number of abdominal wounds + (1|experiment)

**Table S1. Number of wild female *D. melanogaster* and *D. simulans* collected (species combined).** Total numbers collected are given according to whether they did, or did not, produce offspring when brought into the lab, the collection site (D = Domäne Dahlem; L = Obsthof Lindicke and G = SL Gartenbau) and the season (ES = early summer, LS = late summer, A = autumn).

| *Females produced offspring* | *Collection site* | *Season* | | |
|---|---|---|---|---|
| | | *ES* | *LS* | *A* |
| Yes | D | 48 | 53 | 3 |
| | G | 37 | 33 | 60 |
| | L | 87 | 64 | 47 |
| No | D | 8 | 5 | 5 |
| | G | 5 | 9 | 10 |
| | L | 12 | 19 | 12 |

### Supplementary tables

**Table S2: Experimental design for experiment 1.** The table shows the treatment groups and their abbreviated names as used in the figures and the main text, mating events, and genital wound examination times. The subscript "d" indicates days, and they are counted in days post adult eclosion. The + indicate when a mating event occurred or when genital wounds were examined. *Females in the virgin groups were not offered a male.

| Treatment (days post-eclosion) | Abbreviated treatment group name | Day 5 1st mating opportunity* | Day 6 Genital wound examination | Day 7 2nd mating opportunity* | Day 8 Genital wound examination |
|---|---|---|---|---|---|
| Virgin (day 5, n = 57) | $0_{d5}$ | + | + | | |
| Virgin (day 7, n = 57) | $0_{d7}$ | | | + | + |
| Single mating (day 5, n = 61) | $1_{d5}$ | + | + | | |
| Double mating (days 5 and 7, n = 41) | $2_{d5\ d7}$ | + | | + | + |

**Table S3: Experimental design for experiment 2**. The table shows the the treatment groups and their abbreviated names as used in the figures and the main text, mating events, and timing of abdominal and genital wound examinations. The subscript "d" indicates days, and they are counted in days post adult eclosion. The + indicate when an abdominal or genital wound examination was conducted or when a mating event occurred. The $n_{gw}$ refers to the number of flies used for genital wounding assay and $n_{aw}$ refers to the number of flies used for abdominal wounding assay. *Females in the virgin groups were not offered a male. Grey shaded rows indicate treatment groups that were used in experiment 2 but not in experiment 1.

| Treatment (days post-eclosion) | Abbreviated treatment group name | Day 4 Abdominal wound examination | Day 5 1st mating opportunity* | Day 6 Abdominal wound examination | Day 6 Genital wound examination | Day 7 2nd mating opportunity* | Day 8 Abdominal wound examination | Day 8 Genital wound examination |
|---|---|---|---|---|---|---|---|---|
| Virgin (day 5, $n_{gw}$ = 30, $n_{aw}$ = 30) | $0_{d5}$ | + | + | + | | | | |
| Virgin (day 7, $n_{gw}$ = 30, $n_{aw}$ = 30) | $0_{d7}$ | | | + | | + | + | |
| Single mating (day 5, $n_{gw}$ = 39, $n_{aw}$ = 70) | $1_{d5}$ | + | + | + | + | | | |
| Single mating (day 7, $n_{gw}$ = 49, $n_{aw}$ = 83) | $1_{d7}$ | | | + | | + | + | + |
| Single mating, no second mating (day 5, no day 7, $n_{gw}$ = 45, $n_{aw}$ = 46) | $1_{d5,\ no\ d7}$ | | + | | | + | + | + |
| Double mating (days 5 and 7, $n_{gw}$ = 23, $n_{aw}$ = 33) | $2_{d5\ d7}$ | | + | | | + | + | + |

**Table S4. The effects of the mating number and fly population on the amount of time it took to start mating (latency) and the duration of mating.** Statistically significant p-values are in bold font.

| *Experiment* | *Response variable* | *Tested effect* | *df* | *Chisq* | *p* |
|---|---|---|---|---|---|
| 1 | Log Mating latency (mins) | First or second mating | 1 | 21.26 | **< 0.0001** |
| | | Population | 1 | 4.60 | **0.032** |
| | | First or second mating × Population | 1 | 0.56 | 0.46 |
| | Mating duration (mins) | First or second mating | 1 | 4.40 | **0.036** |
| | | Population | 1 | 1.72 | 0.19 |
| | | First or second mating × Population | 1 | 0.16 | 0.69 |
| 2 | Log Mating latency (mins) | First or second mating | 1 | 10.18 | **0.0014** |
| | Mating duration (mins) | First or second mating | 1 | 0.21 | 0.64 |

**Table S5. The effects of mating latency and duration of the first mating on wound number and area.** The same models were run with and without the females that did not produce larvae after the first mating. Sample sizes are given in the table.

| *Experiment* | *Response variable* | *Tested effect* | *df* | *Chisq* | *p* |
|---|---|---|---|---|---|
| 1 | Wound number (n = 61; only females producing larvae) | Mating latency | 1 | 1.62 | 0.20 |
| | | Mating duration | 1 | 1.46 | 0.23 |
| | Wound number (n = 66; all females) | Mating latency | 1 | 2.73 | 0.098 |
| | | Mating duration | 1 | 2.04 | 0.15 |
| | Log Wound area (n = 61; only females producing larvae) | Mating latency | 1 | 0.0017 | 0.97 |
| | | Mating duration | 1 | 0.29 | 0.59 |
| | Log Wound area (n = 67; all females) | Mating latency | 1 | 0.27 | 0.60 |
| | | Mating duration | 1 | 0.16 | 0.69 |
| 2 | Wound number (n = 132) | Mating latency | 1 | 2.03 | 0.15 |
| | | Mating duration | 1 | 0.03 | 0.86 |
| | Log Wound area (n = 132) | Mating latency | 1 | 3.17 | 0.075 |
| | | Mating duration | 1 | 2.79 | 0.095 |

**Table S6. Post-hoc comparisons examining the effects of the number of matings on the presence of genital wounds.** Statistically significant p-values are in bold font.

| *Experiment* | *Response variable* | *Comparisons* | *z ratio* | *p* |
|---|---|---|---|---|
| 2 | Number of wounds | $1_{d5} - 1_{d7}$ | 0.24 | 1.00 |
| | | $1_{d5} - 2_{d57}$ | -3.51 | **0.0025** |
| | | $1_{d5} - 1_{d5,\ no\ d7}$ | -1.66 | 0.34 |
| | | $1_{d7} - 2_{d57}$ | -4.06 | **0.0003** |
| | | $1_{d7} - 1_{d5,\ no\ d7}$ | -2.07 | 0.16 |
| | | $2_{d57} - 1_{d5,\ no\ d7}$ | 2.30 | 0.10 |
| | Wound area | $1_{d5} - 1_{d7}$ | 2.29 | 0.10 |
| | | $1_{d5} - 2_{d57}$ | -3.07 | **0.0135** |
| | | $1_{d5} - 1_{d5,\ no\ d7}$ | -2.69 | **0.0392** |
| | | $1_{d7} - 2_{d57}$ | -5.15 | **< 0.0001** |
| | | $1_{d7} - 1_{d5,\ no\ d7}$ | -5.23 | **< 0.0001** |
| | | $2_{d57} - 1_{d5,\ no\ d7}$ | 0.86 | 0.82 |

**Table S7. The effects of the number of matings and the population on the presence of abdominal wounds.** Mating treatment has three levels: virgin, one mating or two matings. Statistically significant p-values are in bold font.

| *Experiment* | *Response variable* | *Tested effect* | *df* | *Chisq* | *p* |
|---|---|---|---|---|---|
| 1 | Abdominal wounds (n = 180) | Mating treatment | 2 | 72.29 | < **0.0001** |
| | | Population | 1 | 0.44 | 0.50 |
| | Abdominal wounds (n = 170; only mated females producing larvae) | Mating treatment | 2 | 69.10 | < **0.0001** |
| | | Population | 1 | 0.048 | 0.83 |
| 2 | Abdominal wounds (n = 165) | Mating treatment | 3 | 52.15 | < **0.0001** |

**Table S8. Post-hoc comparisons examining the effects of the number of matings and the population on the presence of genital wounds**. Statistically significant p-values are in bold font.

| *Experiment* | *Comparisons* | *z ratio* | *p* |
|---|---|---|---|
| 1 | $0_{d5}$ - $1_{d5}$ | -7.53 | **< 0.0001** |
| | $0_{d5}$ - $2_{d57}$ | -5.31 | **< 0.0001** |
| | $1_{d5}$ - $2_{d57}$ | -0.20 | 0.98 |
| 2 | $1_{d5} – 1_{d7}$ | -0.83 | 0.84 |
| | $1_{d5} – 1_{d5,\ no\ d7}$ | -4.60 | **< 0.0001** |
| | $1_{d5} – 2_{d57}$ | -6.09 | **< 0.0001** |
| | $1_{d7} – 1_{d5,\ no\ d7}$ | -3.91 | **0.0005** |
| | $1_{d7} – 2_{d57}$ | -5.54 | **< 0.0001** |
| | $1_{d5,\ no\ d7} – 2_{d57}$ | -2.23 | 0.11 |

**Table S9. The relationship between the number of abdominal wounds and both the number of genital wounds and the area of genital wounds.** Statistically significant p-values are in bold font.

| *Experiment* | *Response variable* | *Tested affect* | *df* | *Chisq* | *P* |
|---|---|---|---|---|---|
| 2 | Number of genital wounds | Number of abdominal wounds | 1 | 0.29 | 0.59 |
| | | Treatment | 3 | 2.51 | 0.47 |
| | | Number of abdominal wounds × treatment | 3 | 0.19 | 0.98 |
| | Log Wound area | Number of abdominal wounds | 1 | 0.31 | 0.58 |
| | | Treatment | 3 | 22.29 | **< 0.0001** |
| | | Number of abdominal wounds × treatment | 3 | 8.65 | **0.0343** |

### Supplementary figures

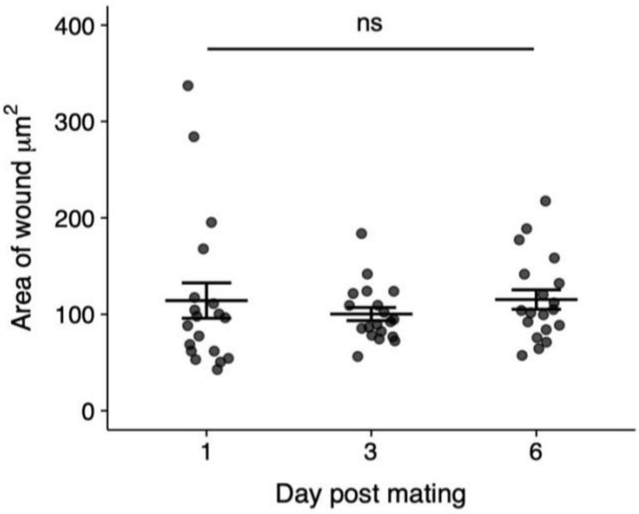


**Figure S1. Area of wounds from females sampled 1, 3 or 6 days after mating**. A preliminary experiment was carried out where females were anaesthatised with $CO_2$ and sacrificed by being placed into ethanol either 1-, 3-, or 6-days post mating. On the seventh day post mating all females were dissected in a random order and blind with respect to treatment. All other methods were the same as described in the materials and methods. The area of wounding did not differ according to the day after mating at which the females were sacrificed ($F_{2,54} = 0.44$, $p = 0.65$).

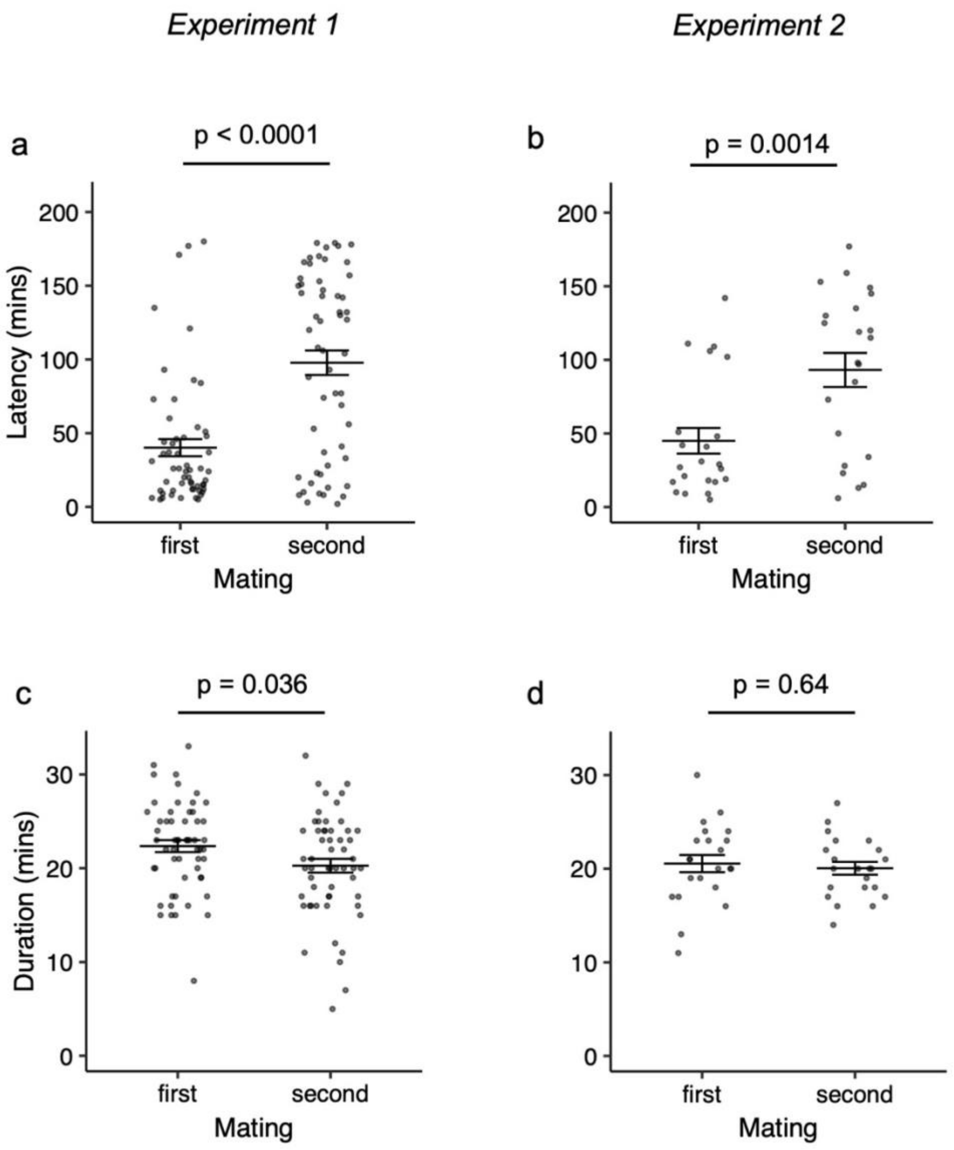


**Figure S2. Latency and duration of first and second matings.** Latency, that is the time until copulation was initiated, for (**a**) experiment 1 and (**b**) experiment 2. Copulation duration for (**c**) experiment 1 and (**d**) experiment 2. Only females who mated twice were included in the data, i.e., the data are paired, whereby females assayed for the first mating were also assayed for the second mating (experiment 1, n = 56 females; experiment 2, n = 22). Means and standard errors are shown.

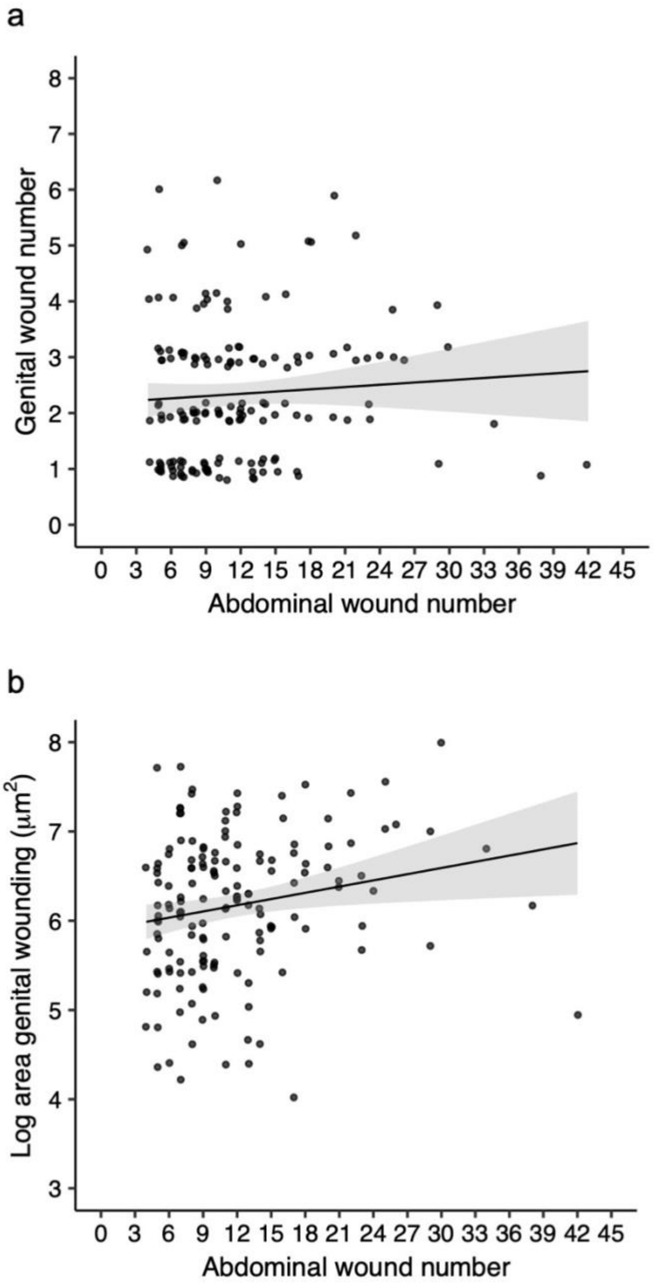


**Figure S3. Experiment 2: Correlations between the two types of copulatory wounding.** (**a**) Relationship between abdominal wound number and genital wound number. (**b**) Relationship between abdominal wound number and genital wound area. Neither of the relationships are statistically significant.
